# Crystal structure of the potato leafroll virus coat protein and implications for viral assembly

**DOI:** 10.1101/2021.08.03.454973

**Authors:** Myfanwy C. Adams, Carl J. Schiltz, Michelle L. Heck, Joshua S. Chappie

## Abstract

Luteoviruses, poleroviruses, and enamoviruses are insect-transmitted, agricultural pathogens that infect a wide array of staple food crops. Previous cryo-electron microscopy studies of virus-like particles indicate that luteovirid viral capsids are built from a structural coat protein that organizes with T=3 icosahedral symmetry. Here we present the crystal structure of a truncated version of the coat protein monomer from potato leafroll virus at 1.57-Å resolution. In the crystal lattice, monomers pack into flat sheets that preserve the two-fold and three-fold axes of icosahedral symmetry and show minimal structural deviations when compared to the full-length subunits of the assembled virus-like particle. These observations have important implications in viral assembly and maturation, suggesting that the CP N-terminus and its interactions with RNA serve as a key driver for generating capsid curvature.

## 1. Introduction

Viral capsids exhibiting icosahedral symmetry are constructed with 60 identical subunits arranged around 12 vertices with 5-fold rotational symmetry, 20 triangular faces with 3-fold symmetry, and 30 edges with 2-fold symmetry (Sevvana et al., 2021). Icosahedral viruses containing more than 60 subunits follow the principles of quasi-equivalence (Caspar and Klug, 1962), wherein identical subunits form both hexamer and pentamer morphological units that serve as the building blocks for the quasi-equivalent viral particle (Johnson, 1996). The triangulation number *T* (*T*=*h*^2^+*hk*+*k*^2^, where *h* and *k* are integers) relates how pentamers can be inserted into a hexagonal lattice to produce an enclosed icosahedron and defines the discrete geometric subdivisions of the triangular face needed to accommodate an increased number of subunits (Rossmann, 2013). By these rules, capsids containing 60*T* subunits will arrange as 12 pentamers and 10(*T*-1) hexamers with the underlying icosahedral asymmetric unit containing *T* subunits that are quasi-equivalent to each other (Johnson and Speir, 1997; Prasad and Schmid, 2011). A T=3 virus therefore contains 180 total subunits (60 x 3) with three quasi-equivalent monomers forming each triangular face of the icosahedron.

Luteoviruses, poleroviruses, and enamoviruses (collectively referred to as ‘luteovirids’) are economically important agricultural pathogens that devastate many staple food crops (Ali et al., 2014). Luteovirids replicate exclusively in plant hosts and are phloem-limited, relying on particular species of phloem-feeding insects for efficient transmission in a circulative manner that involves movement across and within specific insect tissues (Gray et al., 2014; Gray and Gildow, 2003; Whitfield et al., 2015). These viruses contain a single-stranded, positive sense RNA genome that encodes two major structural proteins (Miller et al., 1995): a coat protein (CP) and a minor readthrough protein (RTP) that is translated via stochastic ribosomal readthrough of the CP stop codon (Brault et al., 1995; Cheng et al., 1994; Dong et al., 1998; Filichkin et al., 1994) (Figure 1A). The readthrough domain (RTD) that is added to the CP is not required for virion assembly or infection (Filichkin et al., 1994; Reutenauer et al., 1993) but plays a critical role in viral uptake and transmission by aphids (Bruyère et al., 1997; Chay et al., 1996; Gildow et al., 2000; Peter et al., 2008; Reinbold et al., 2001) and has been implicated in the long-range movement (Boissinot et al., 2014) and phloem-limitation (Peter et al., 2009) of the virus in plants. Cryo-electron microscopy (cryo-EM) characterization of polerovirus virus-like particles (VLPs) lacking the RTD confirmed that luteovirid capsids assemble with T=3 icosahedral symmetry and identified the key interfaces and contacts between CP subunits that maintain the structural integrity of the virion (Byrne et al., 2019). Here we present the crystal structure of a truncated version of the coat protein monomer from potato leafroll virus (PLRV^68-208^) at 1.57-Å resolution. In the crystal lattice, PLRV^68-208^ packs into flat sheets that are stabilized exclusively by hexameric interactions. This organization preserves the two-fold and three-fold axes of icosahedral symmetry but lacks the curvature necessary to generate enclosed VLPs. Structural comparison between PLRV^68-208^ and VLP subunits shows minimal structural differences across the CP monomer core, implicating the truncated N-terminus and its interactions with RNA as an important driver of viral assembly and maturation.

**Figure 1.**
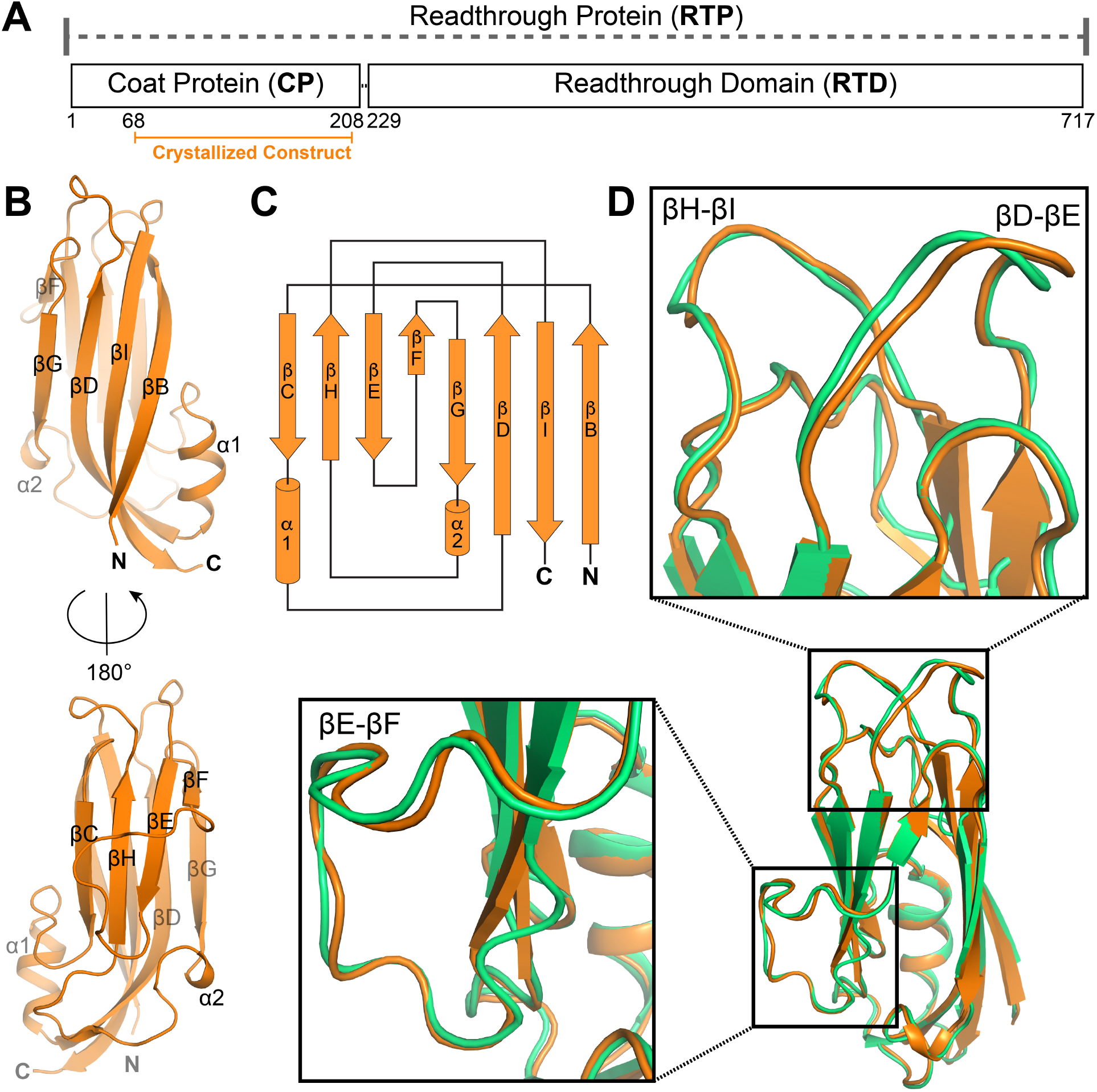
Structure of truncated PLRV CP monomer. A. Domain architecture of PLRV readthrough protein (RTP). Boundaries of the coat protein (CP) and readthrough domain (RTD) are labeled and numbered. Orange denotes purified fragment (PLRV^68-208^) used for crystallographic studies. B,C. Structure (B) and topology (C) of PLRV^68-208^ monomer. D. Superposition of PLRV^68-208^ chain A (orange) with full-length CP monomer derived from the cryo-EM reconstruction of the PLRV virus-like particle (green, PDB: 6SCO, chain A). Black boxes highlight zoomed views of conformational differences.

## 2. Materials and methods

### 2.1 Cloning, expression, and purification of PLRV^68-208^

DNA encoding PLRV minimal capsid domain (residues 68-208) was cloned from an infective PLRV isolate and inserted into pCAV4, a modified T7 expression vector that introduced an N-terminal 6xHis-NusA tag followed by a HRV 3C protease site upstream of the inserted sequence. Constructs were transformed into BL21(DE3) cells, grown at 37°C in Terrific Broth to an OD_600_ of 0.7-0.9, and then induced with 0.3 mM IPTG overnight at 19°C. Cells were pelleted and resuspended in nickel loading buffer (20 mM HEPES pH 7.5, 500 mM NaCl, 30 mM imidazole, 5% glycerol (v:v), and 5 mM β-mercaptoethanol) supplemented with 0.5 mg DNAse, 10 mM MgCl_2_, 1 mM PMSF, and a Roche complete protease inhibitor cocktail tablet. Lysozyme was added to a concentration of 1 mg/ml and the mixture was incubated for 10 minutes rocking at 4°C. Cells were disrupted by sonication and the lysate was cleared via centrifugation at 13 000 rpm (19 685 g) for 30 minutes at 4°C. The supernatant was filtered, loaded onto a 5 ml HiTrap chelating column charged with NiSO4, and then washed with nickel loading buffer. PLRV^68-208^ was eluted by an imidazole gradient from 30 mM to 500 mM. HRV 3C protease was added to the pooled fractions, which were subsequently dialyzed overnight at 4°C against S buffer (20 mM HEPES pH 7.5, 50 mM NaCl, 1 mM EDTA, 5% glycerol (v:v), and 1 mM DTT). Significant precipitation occurred during dialysis; however, uncut NusA-PLRV^68-208^ constituted the vast majority of the insoluble fraction. The clarified protein sample was applied to a 5 ml HiTrap S column equilibrated with S buffer. The sample was washed with S buffer and eluted with a NaCl gradient from 50 mM to 500 mM over 12 column volumes. Peak S column fractions were pooled, concentrated, and injected onto a Superdex 75 10/300 GL sizing column equilibrated in 20 mM HEPES pH 7.5, 150 mM KCl, and 1 mM DTT. The protein was concentrated to 10-20 mg/ml, flash frozen in liquid nitrogen, and stored at −80°C.

### 2.2 Crystallization, X-ray data collection, and structure determination

PLRV^68-208^ (6-10 mg/mL) was mixed in a 1:1 ratio with 0.1 M Bis-Tris Propane pH 7.7, 26% PEG 3350, and 0.1 M NaBr. The highest quality crystals were obtained using sitting drop vapour diffusion with a drop size of 2 uL and reservoir volume of 65 uL at 20°C. Samples were cryoprotected by transferring the crystal directly to Parabar 10312 (Hampton Research) prior to freezing in liquid nitrogen. Crystals were of the space group P1 with unit cell dimensions a = 36.86 Å, b = 67.49 Å, c = 75.88 Å and α = 79.66°, β = 76.21°, γ = 81.64°. Data were collected remotely on the tuneable NE-CAT 24-ID-C beamline at the Advanced Photon Source at the selenium edge energy at 12.663 kEv (Table 1). Data were integrated and scaled using XDS (Kabsch, 2010) and AIMLESS (Evans and Murshudov, 2013) via the NE-CAT RAPD pipeline. The structure was solved by molecular replacement using PHASER (McCoy and Read, 2010) and a monomer from the cryo-EM structure of the PLRV VLP (PDB: 6SCO) as the search model. Further model building and refinement was carried out manually in COOT (Emsley et al., 2010) and PHENIX (Liebschner et al., 2019), respectively. The final model was refined to 1.57-Å resolution with R_work_/R_free_ = 0.1792/0.2281 (Table 1) and contained six identical molecules in the asymmetric unit: chains A-F, 68-208. All structural models were rendered with Pymol (Schrodinger).

**Table 1.** X-ray data collection and refinement statistics for PLRV^68-208^.

| **Data collection** |  |
| --- | --- |
| PDB code | 7RLM |
| X-ray Source | NECAT 24-ID-C |
| Wavelength (Å) | 0.9791 |
| Space group | P1 |
| Unit cell | $a = 36.86, b = 67.48, c = 75.89$ Å<br>$\alpha = 79.67^\circ, \beta = 76.22^\circ, \gamma = 81.67^\circ$ |
| Resolution, Å<sup>a</sup> | 72.86 – 1.504 (1.558 – 1.504) |
| No. measured reflections<sup>a</sup> | 323209 (24594) |
| No. unique reflections<sup>a</sup> | 177750 (14912) |
| Completeness (%)<sup>a</sup> | 89.1 (46.8) |
| Multiplicity<sup>a</sup> | 3.3 (1.5) |
| R<sub>merge</sub><sup>a</sup> | 0.072 (0.868) |
| Mean I/σ<sub>I</sub><sup>a</sup> | 8.6 (0.5) |
| CC<sub>1/2</sub><sup>a</sup> | 0.997 (0.315) |
| **Refinement** |  |
| Resolution (Å) | 72.86-1.57 |
| No. relections | 90356 (8750) |
| R<sub>work</sub>/R<sub>free</sub> | 0.1792 / 0.2281 |
| RMSD |  |
| Bond lengths (Å) | 0.006 |
| Bond angles (°) | 0.823 |
| Ramachandran plot |  |
| Favored (%) | 97.96 |
| Allowed (%) | 2.04 |
| Outliers (%) | 0.00 |
| Average B-Factor | 26.14 |
| Clash score | 5.53 |
| No. Atoms |  |
| Macromolecule | 6643 |
| Solvent | 686 |
| Other ions if present? | 1 |

<sup>a</sup> Parentheses indicate values for highest resolution shell

## 3. Results and discussion

### 3.1 Structural fold of truncated PLRV^68-208^

PLRV is phloem limited, making it difficult to obtain sufficient quantities of mature, infectious virions from natural sources for comprehensive structural studies. We instead designed a soluble, PLRV CP construct (PLRV^68-208^) that could be expressed in *E. coli* and purified in milligram quantities. The domain boundaries of this construct were chosen based on disorder prediction (Erdős and Dosztányi, 2020), truncating the flexible, arginine-rich N-terminus (residues 1-67) that ordinarily interacts with viral RNA (Figure 1A). Purified PLRV^68-208^ exists as a monodispersed, unassembled monomer in solution and produced small, needle-like crystals that routinely diffracted to ~1-2 Å. Crystals were of the space group P1 with six molecules in the asymmetric unit. The structure was solved by molecular replacement using a monomer from the PLRV VLP cryo-EM structure (PDB: 6SCO) as a search model. The final model was refined to 1.57-Å resolution with R_work_ and R_free_ values of 0.1792/0.2281 (Table 1).

The truncated PLRV^68-208^ monomer adopts a jelly-roll fold composed of two antiparallel beta sheets – topologically ordered βB-βI-βD-βG and βF-βE-βH-βC (Rossmann and Johnson, 1989) – that sandwich together and are flanked by two a-helices (Figure 1B-C). Superposition shows good overall structural agreement between PLRV^68-208^ and the full-length CP monomer that make up the asymmetric unit of the PLRV VLP cryo-EM structure (Figure 1D), with overall RMSDs across backbone atoms on the order ~0.5 Å when each chain is compared (Table 2). The main structural variations occur in the positions of the βD-βE, βE-βF, and βH-βI loops (Figure 1D). This indicates that the core CP fold remains largely unaffected by the underlying lattice organization.

**Table 2.** Overall RMSD values calculated from superposition of PLRV CP subunits.

|  |  | PLRV<sup>68-208</sup> |  |  |  |  |  | PLRV VLP |  |  |
| --- | --- | --- | --- | --- | --- | --- | --- | --- | --- | --- |
|  |  | A | B | C | D | E | F | A | B | C |
| PLRV<sub>68-208</sub> | A | - | 0.296 | 0.247 | 0.278 | 0.278 | 0.32 | 0.525 | 0.525 | 0.525 |
|  | B | 0.296 | - | 0.320 | 0.284 | 0.219 | 0.313 | 0.552 | 0.552 | 0.552 |
|  | C | 0.247 | 0.320 | - | 0.268 | 0.262 | 0.343 | 0.555 | 0.555 | 0.555 |
|  | D | 0.278 | 0.284 | 0.268 | - | 0.265 | 0.355 | 0.561 | 0.561 | 0.561 |
|  | E | 0.278 | 0.219 | 0.262 | 0.265 | - | 0.324 | 0.560 | 0.560 | 0.560 |
|  | F | 0.320 | 0.313 | 0.343 | 0.355 | 0.324 | - | 0.521 | 0.521 | 0.521 |
| PLRV VLP | A | 0.525 | 0.552 | 0.555 | 0.561 | 0.560 | 0.521 | - | 0.001 | 0.001 |
|  | B | 0.525 | 0.552 | 0.555 | 0.561 | 0.560 | 0.521 | 0.001 | - | 0.001 |
|  | C | 0.525 | 0.552 | 0.555 | 0.561 | 0.560 | 0.521 | 0.001 | 0.001 | - |

### 3.2 Crystal packing demonstrates innate preference for elements of icosahedral symmetry

PLRV^68-208^ monomers pack in layers of flat sheets that stack on top of one another within the crystal lattice (Figure 2A). The crystallographic asymmetric unit spans two of these layers, with three molecules localized to each plane in different orientations (Figure 2A-B). Within a single sheet, the protein monomers form an ordered array that contains two-fold, three-fold, and six-fold non-crystallographic symmetry (Figure 2C). This organization recapitulates the subunit interfaces found within the assembled capsid of PLRV VLPs as well as the two-fold and three-fold icosahedral symmetry axes (Figure 2D). VLP assembly with a five-fold axis in place of a six-fold axis imposes constraints that induce significant curvature, allowing closure of the icosahedron (Figure 2E). Direct comparison of the planar (Figure 2C) and icosahedral (Figure 2D) asymmetric units shows that a slight downward rotation of the B and C subunits and ~9° upward tilt of the A subunit accompanies the underlying symmetry transition from flat sheet to mature capsid (Figure 2F).

**Figure 2.**
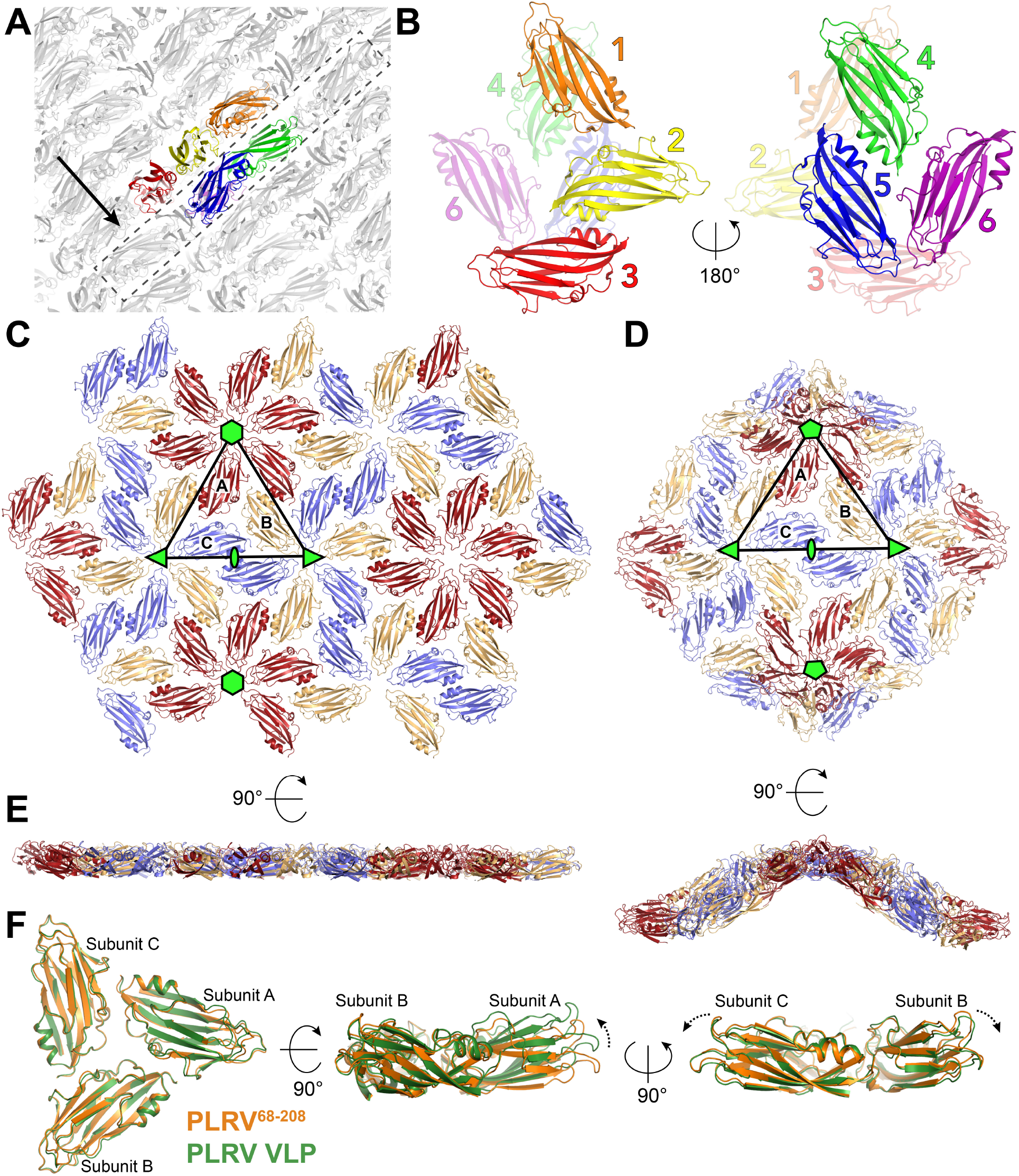
Crystal packing of PLRV^68-208^. A. CP monomers (gray) pack into planar sheets (dashed box) that are stacked on top of each other within the PLRV^68-208^ crystal lattice. Colored monomers indicate relative position of the asymmetric unit, which spans two sequential layers. Black arrow denotes orientation of the view in panel C. B. Spatial distribution of the six monomers in the asymmetric unit. C. Two-dimensional packing of PLRV^68-208^ monomers in the crystal lattice. Non-crystallographic two-fold (green ellipse), three-fold (green triangles), and six-fold (green hexagons) symmetry axes are marked. Monomers that comprise the planar asymmetric unit (black triangular outline) are colored red, light orange, and slate and labeled ‘A’, ‘B’, and ‘C’, respectively. D. T=3 icosahedral symmetry of PLRV VLPs (PDB: 6SCO). Individual subunits that constitute the icosahedral asymmetric unit (black triangular outline) are colored red, light orange, and slate and labeled ‘A’, ‘B’, and ‘C’, respectively. Two-fold (green ellipse), three-fold (green triangles), and five-fold (green pentagons) icosahedral symmetry axes are marked. E. Relative curvature differences between of PLRV^68-208^ crystal lattice (left) and PLRV VLPs (right). F. Structural superposition of the planar (orange) and icosahedral green) asymmetric units from C and D. Dashed black arrows indicate relative movements that propagate curvature in the viral capsid.

### 3.3 Implications for viral capsid assembly

In developing the conceptual framework of quasi-equivalence, Caspar and Klug argued that (i) the inter-subunit bonding energy serves as the driving force for assembly, (ii) a closed shell maximizes the number of bonds between subunits, and (iii) introduction of pentamers is required to achieve the natural curvature needed for shell closure, even if hexamers are intrinsically stable enough to be formed preferentially (Caspar and Klug, 1962). The distinct PLRV CP assemblies captured by X-ray crystallography and cryo-EM exemplify these principles. PLRV^68-208^ exclusively forms hexameric interactions within the crystal lattice and thus remains organized in planar sheets. In contrast, assembly of full-length CP monomers *in planta* efficiently incorporates pentamers, leading to stable VLPs. These structural changes occur with minimal distortions to the overall fold and without significant perturbations to the primary intersubunit interactions, providing experimental support for Caspar and Klug’s theoretical descriptions.

The PLRV VLPs used for cryo-EM studies were generated by transiently expressing the full-length CP in *Nicotiana benthamiana* following agroinfiltration (Byrne et al., 2019). The resulting particles contain an interior amorphous density that likely consists of the N-terminal domains and/or packaged RNA (Byrne et al., 2019). Our attempts to express the full-length PLRV CP in *E. coli* yielded only unfolded aggregates. Luteovirid VLPs have also been produced using baculovirus expression systems in insect cells (Gildow et al., 2000; Lamb et al., 1996; Sivakumar et al., 2009; Tian et al., 1995). For PLRV, the native CP was insoluble in Sf9 cells but could be coaxed to form VLPs when an extended histidine tag was added at the N-terminus (Gildow et al., 2000; Lamb et al., 1996). His-tagged pea enation mosaic virus 1 (PEMV) CP constructs expressed in Sf21 cells formed VLPs that were heterogeneous in size and contained baculoviral mRNA (Sivakumar et al., 2009). Non-tagged PEMV VLPs could not be purified (Sivakumar et al., 2009), however, mirroring the behavior of baculovirus-derived beet western yellow virus VLPs (Tian et al., 1995). These observations suggest that accessory factors in host plants likely play a critical role in the proper assembly of luteovirid capsids from native CP subunits. Cross-linking coupled to high resolution mass spectrometry previously identified plant chaperones that interact with the PLRV CP and may play a role in this process (DeBlasio et al., 2016).

The T=3 icosahedral symmetry of PLRV capsids dictates that the A, B, and C subunits within the icosahedral asymmetric unit (Figure 2D) are quasi-equivalent despite deriving from identical gene products. In many T=3 viruses, the flexible N-terminal portion of the capsid protein mediates conformational differences that facilitate non-equivalent interactions (Prasad and Schmid, 2011), be it through order-to-disorder transitions (Abad-Zapatero et al., 1980; Harrison et al., 1978; Hogle et al., 1986; Prasad et al., 1999), direct interactions with genomic RNA (Fisher and Johnson, 1993; Makino et al., 2013), or residue-specific conformational changes (Chen et al., 2006; Ossiboff et al., 2010). Mutagenesis and/or deletion of this N-terminal region often disrupts capsid assembly (Bertolotti-Ciarlet et al., 2002; Dong et al., 1998; Wikoff and Johnson, 1999), underscoring its importance in virion stability and icosahedral quasi-equivalent switching. We speculate that the CP N-terminus plays a similar regulatory role in PLRV based on PLRV^68-208,^s inability to form closed capsids and the non-symmetric cryo-EM density present in the interior of the PLRV VLPs, which would be occupied by the residues removed from the truncated CP construct and/or scavenged RNA that associates with them.

### 3.4 Concluding remarks

Understanding viral assembly is critical for the development of new strategies to mitigate luteovirid transmission and subsequent crop loss. Our work shows that crystallography is a viable approach to elucidate the structural geometry and symmetry interactions of other luteovirids when the formation of VLPs and/or isolation of stable, mature virions is not possible. The PLRV^68-208^ crystal structure not only affords a high-resolution snapshot of the PLRV CP monomer but also shows a potential intermediate stage of capsid folding. Future *in vitro* studies examining CP assembly and the effects of viral RNA will be necessary to define the exact mechanism and kinetics of PLRV maturation. The PLRV CP assemblies described and compared in this work will help in modelling these pathways.

## 4. Accession numbers

The atomic coordinates and structure factors for the truncated potato leafroll virus coat protein (residues 68-208) have been deposited in the Protein Data Bank (http://www.rcsb.org) under the PDB code 7RLM.

## CRediT authorship contribution statement

**Myfanwy C. Adams:** Conceptualization, Methodology, Validation, Investigation, Writing - original draft, Visualization. **Carl J. Schiltz:** Conceptualization, Methodology, Validation, Investigation, Visualization. **Michelle L. Heck:** Conceptualization, Methodology, Validation, Investigation, Writing - original draft, Supervision, Funding acquisition. **Joshua S. Chappie:** Conceptualization, Methodology, Validation, Investigation, Writing - original draft, Visualization, Supervision, Funding acquisition.

## Acknowledgements

We thank the Northeastern Collaborative Access Team (NE-CAT) beamline staff at the Advanced Photon Source (APS) for assistance with remote X-ray data collection. This work was supported by USDA Agricultural Research grant 2019-05200 (to M.L.H. and J.S.C) and is based upon research conducted at the Northeastern Collaborative Access Team (NE-CAT) beamlines under GUP-62147 (PI: J.S.C.). NE-CAT is funded by the National Institute of General Medical Sciences from the National Institutes of Health (P30 GM124165). The Pilatus 6M detector on 24-ID-C beam line is funded by a NIH-ORIP HEI grant (S10 RR029205). This research used resources of the Advanced Photon Source, a U.S. Department of Energy (DOE) Office of Science User Facility operated for the DOE Office of Science by Argonne National Laboratory under Contract No. DE-AC02-06CH11357. J.S.C. is a Meinig Family Investigator in the Life Sciences. M.C.A. is supported by a NIFA predoctoral fellowship (2020-67034-31750). USDA is an equal opportunity provider and employer.

